# Cryptic microgeographic variation in responses of larval Atlantic cod to warmer temperatures

**DOI:** 10.1101/2021.02.03.429645

**Authors:** Rebekah A. Oomen, Elisabeth Juliussen, Esben M. Olsen, Halvor Knutsen, Sissel Jentoft, Jeffrey A. Hutchings

**Affiliations:** Department of Biology, Dalhousie University, Halifax, Nova Scotia, Canada; Centre for Ecological and Evolutionary Synthesis (CEES), Department of Biosciences, University of Oslo, Oslo, Norway; Centre for Coastal Research□(CCR), University of Agder, Kristiansand, Norway; Institute of Marine Research, Flødevigen Marine Research Station, His, Norway; County Governor in Agder, Arendal, Norway

**Keywords:** adaptation, climate change, common garden experiment, fisheries management, phenotypic plasticity, temperature

## Abstract

Although temperature is known to drive species dynamics and distributions, our understanding of the extent to which thermal plasticity varies within species is poor. Differences in plasticity can arise through local adaptation to heterogeneous environments, hybridization, and the release of cryptic genetic variation in novel environments. Here, wild Atlantic cod (*Gadus morhua*) from contrasting environments inside and outside of a fjord system in southern Norway spawned freely in a semi-natural laboratory environment, generating pure crosses and reciprocal hybrids. A common-garden rearing experiment of the larvae at 6°C, 9.5°C, and 13°C revealed cryptic genetic variation in thermal responses of growth and survival at warmer temperatures. Variation in growth plasticity was greatest from 9.5°C to 13°C, the latter of which exceeds temperatures currently typical of larvae in their native environments. In contrast to our prediction of intermediate hybrid responses consistent with additive genetic effects, one reciprocal hybrid cross showed a 4% increase in size at the highest temperature, whereas most crosses exhibited 4-12% reductions in size. All crosses experienced severe (76-93%) reductions in survival from 9.5°C to 13°C. Variation in survival plasticity suggests a genetically variable basis for the severity with which survival declines with increasing temperature and the potential for an adaptive response to warming. Notably, we demonstrate the potential for hybridization between coexisting ‘fjord’ and ‘North Sea’ ecotypes that naturally inhabit the inner and outer fjord environments at contrasting frequencies. Yet, ecotype explained a minor (3-10%) component of growth reaction norm variation, suggesting it is insufficient for describing important biological variation. Current broad-scale management and lack of coastal monitoring impede the development of strategies to maintain the potential for adaptation to warming temperatures in systems with such phenotypic complexity resulting from cryptic genetic variation, coexisting ecotypes, and gene flow.

## Introduction

Adaptive phenotypic plasticity is a primary mechanism by which species respond to environmental change, by serving as a short-term buffer against environmental variability (Canale & Henry, 2010) and potentially facilitating evolutionary adaptation to new environments over multiple generations (Chevin, Lande, & Mace, 2010; Lande, 2009).

Intraspecific variation in adaptive plasticity can arise when it evolves in response to local environmental regimes (e.g. Conover & Present 1990; McCairns & Bernatchez 2010; Baumann & Conover 2011). Non-adaptive (i.e., not shaped *via* adaptation) plasticity in response to novel or extreme environments can also facilitate adaptive evolution by releasing cryptic genetic variation that increases the variance around the mean response and can result in a fitter phenotype by chance (Ghalambor, McKay, Carroll, & Reznick, 2007; Murren et al., 2014). Whether heritable variation in plasticity has arisen adaptively or non-adaptively, species can respond to environmental change through adaptation if sufficient heritable variation exists for selection to act on (Franks & Hoffman, 2012) and adaptation can keep pace with the rate of environmental change (Hoffmann & Sgrò, 2011). Therefore, intraspecific variation in plasticity likely represents differences in the responses of individuals and populations to directional environmental change, such as the forecasted increase in temperature due to the global climate crisis, and the adaptive potential of the species as a whole to endure or colonize new environments.

Understanding the mechanisms responsible for shaping intraspecific variation in environmental responses, and the spatial scales at which differences in plasticity occur, are critical for predicting the persistence of a species in the face of environmental change and managing populations effectively to mitigate the potential for population collapse and biodiversity loss.

The marine environment was traditionally assumed to be genetically homogeneous, due to the high potential for dispersal (Levin, 2006) and limited apparent physical barriers to gene flow (Hilbish, 1996). There is now widespread evidence of adaptation associated with broad-scale spatial variation in selection pressures in species with high connectivity (e.g., Limborg et al. 2012, Pespeni & Palumbi 2013). In marine fishes, divergence in plasticity has been quantified at relatively broad spatial scales across open waters (reviewed by Hutchings 2011; Oomen & Hutchings 2015a). The smallest scale at which genetic variation in plasticity of an adaptive trait (i.e., clearly fitness-linked, in this case survival plasticity) has been found across open waters in a marine fish is ∼200 km, in Atlantic cod (*Gadus morhua*; hereafter, ‘cod’) off the coast of Nova Scotia, Canada (Oomen & Hutchings, 2015b). Although this variation was attributed to temporally variable selection on different spawning components, spatial variability in selection pressures could concurrently drive local adaptation at small scales in the marine environment. For example, highly localized hydrography and the influence of environmental variation on nearby land can generate fine-scale environmental heterogeneity in coastal waters (Albretsen, Aure, Sætre, & Danielssen, 2012; Knutsen et al., 2018). Unlike broad-scale population differentiation, it is not well understood whether the migration-selection balance can skew in favour of local adaptation under the high gene flow potential of shorter distances (Felsenstein, 1976).

Norway has a heterogeneous coast comprising fjords, inlets, islets, skerries and variable depths. As a result, water temperatures, ocean current patterns, salinities and several other biotic and abiotic factors can be spatially structured at scales of less than a few kms (Johannessen & Dahl, 1996; Skjoldal, Hopkins, Erikstad, & Leinaas, 1995). This environment provides opportunities for species to adapt to local conditions at smaller spatial scales than the open ocean. For example, along the Skagerrak coast in southern Norway, Atlantic cod exhibit genetic variation in maturation reaction norms among neighbouring fjords, where the scale of variation is comparable to that of population connectivity inferred from neutral genetic markers (Olsen et al., 2008). Coastal cod in this region is also characterized by fine-scale spatial variation in recruitment and juvenile growth rates, as well as a negative influence of warmer water on these productivity-oriented parameters, raising questions as to the underlying physiological drivers of these processes (Rogers & Stenseth, 2017; Rogers et al., 2011). Thus, the Norwegian Skagerrak coast presents an ideal system for testing for small-scale variation in plasticity of adaptive traits in marine fishes. Key questions are the genetic and phenotypic distinctiveness of local populations inhabiting inner fjord areas, the extent and consequences of their mixing with cod inhabiting outer oceanic regions, and the impacts of temperature on early life stages. These questions are urgent, as Skagerrak coastal cod have remained at historically low abundances since the early 2000s despite implementation of commercial fishing regulations aimed at facilitating recovery (Aglen et al., 2016; ICES, 2018). Delineating spatial scales sufficient for promoting or constraining local adaptation is of critical importance for conservation, as management policies are implemented geographically (ICES, 2018)

Coastal cod in Norway are currently managed as two units, divided north and south of 62° North. South of 62° to the Skagerrak, where our study area is located, is considered part of the North Sea Cod management unit, whereby Norway is responsible for >1000 km of coastline. Significant neutral genetic structure recently documented along the entire Norwegian coast (Dahle et al., 2018) coupled with the discovery of coexisting fjord and North Sea ecotypes in southern coastal waters (Knutsen et al., 2018) have drawn attention to a need for revision of the current management units that accounts for this complex genetic substructure.

The fjord system around Risør represents a central location on the Skagerrak coast that covers an area of 20 km^2^ and is shaped like a ‘U’ (Figure 1). The outer part of the fjord (Østerfjorden, hereby referred to as ‘outer Risør’) contains small islands and is exposed to the Skagerrak and the North Sea. The inner part of the fjord (Sørfjorden, hereby referred to as ‘inner Risør’) is a more sheltered area, characterized by numerous sills with depths ranging from 20 to 40 m (Bergstad, Torstensen, & Bøhle, 1996; Knutsen et al., 2011; Knutsen et al., 2007). These contrasting environments generate fine-scale heterogeneity in thermal regimes that might promote local adaptation in thermal plasticity, especially for the pelagic egg and larval phases that have limited ability to escape undesirable temperatures.

**Figure 1:**
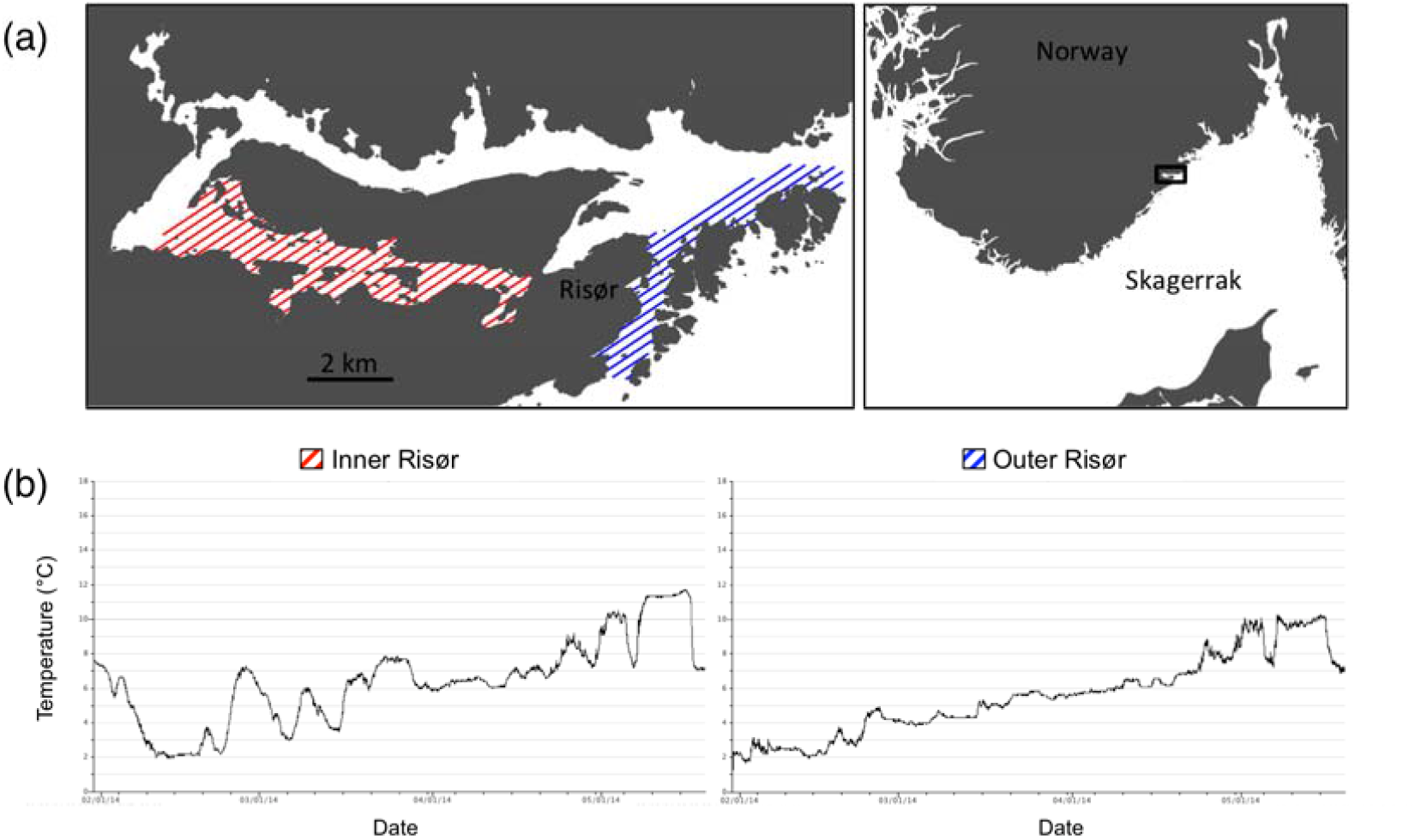
Risør fjord study system on the Norwegian Skagerrak coast. (a) Adult collection locations for inner and outer fjord areas (modified from Kuparinen et al. 2016) and (b) representative temperature profiles during the 2014 spawning season from data loggers positioned at 10 m depth in each study area.

Neutral microsatellite markers have previously revealed a low but significant and temporally stable level of genetic differentiation between juvenile Atlantic cod from inner and outer Risør (Knutsen et al., 2011). It was later discovered, using a much larger set of markers, that this fine-scale spatial divergence is, at least in part, a reflection of the coexistence of two genetically distinct ecotypes that vary in proportion inside and outside the Skagerrak fjords (Barth, Villegas-Ríos, Freitas, Moland, Star, André, et al., 2019; Jorde, Kleiven, et al., 2018; Knutsen et al., 2018; Sodeland et al., in review). The ‘fjord’ ecotype is more common inside the fjord and more likely to be targeted by the local recreational hook-and-line fishery (Jorde, Kleiven, et al., 2018). The ‘North Sea’ ecotype is genetically indistinguishable from the offshore North Sea population. It is not yet known whether the North Sea ecotype disperses through ocean currents as pelagic eggs and larvae from the North Sea to the coastal Skagerrak area, spawns in coastal Skagerrak, or both. However, the stable presence of early life stages and adults in spawning condition suggests they reside in coastal areas and likely recruit to the local population (Jorde, Kleiven, et al., 2018; Jorde, Synnes, Espeland, Sodeland, & Knutsen, 2018; Knutsen et al., 2018; Rogers, Olsen, Knutsen, & Stenseth, 2014).

How these divergent ecotypes persist in sympatry is unknown, as are their ability to hybridize and the phenotypes that distinguish them. Thus far, phenotypic differentiation has only been documented for growth rate and field metabolic rate in wild juveniles, where it might be driven by exposure to different environments earlier in life (Chung et al., 2020; Jørgensen, Neuheimer, Jorde, Knutsen, & Grønkjær, 2020; Knutsen et al., 2018). As phenotypes are the target of natural selection, investigating genetically based variation in phenotypes in coastal cod contributes to filling a major gap in our understanding of whether the genotypic variation observed is biologically and evolutionarily meaningful.

Here, we conduct a spawning and common garden experiment to quantify genetic variation in thermal plasticity for larval growth and survival, to better understand the genetic basis of variable thermal responses in cod and the spatial scale at which such variation exists. The larval stage represents a critical period for survival and development that is especially sensitive to temperature (Pörtner et al., 2008) and for which thermotolerance often most accurately reflects the species climatic boundaries for colonization (Abele, 2012). Specifically, we test for variation in thermal reaction norms associated with fine-scale environmental heterogeneity between inner and outer fjord areas (hereafter, ‘location’) by comparing four genetic crosses: pure inner Risør fjord (I×I), pure outer Risør fjord (O×O), and reciprocal hybrids (I×O and O×I). Using a set of ecotype-associated SNPs (Jorde, Kleiven, et al., 2018), we account for the degree to which these responses are affected by differing proportions of fjord and North Sea ecotypes among crosses. We predicted that 1) genetic variation in plasticity is greater at warmer temperatures not typically experienced during the larval stage, consistent with the release of cryptic genetic variation in extreme environments, and 2) pure crosses would exhibit divergent reaction norms whereas the responses of potential hybrids would be intermediate, consistent with local adaptation and the presence of additive genetic effects.

## Methods

### Temperature data collection

From February to May 2014, 12 temperature loggers were distributed at three depths (5, 10 and 15 m) in four locations; two each in inner and outer Risør (Figure 1). Temperature data were successfully recovered from two depths at one location inside the fjord and from three depths at one location outside the fjord. In February, at the time when spawning initiated in the lab, the mean temperature at 5 metres was 2.95°C (SD±1.21) inside the fjord and 2.78°C (SD±0.84) outside the fjord. Throughout the spawning season, temperatures rose from 2°C to above 8°C at both locations. However, temperatures outside the fjord were more stable and exhibited a smoother increase than temperatures inside the fjord (Figure 1b, Table S1).

### Broodstock

A common-garden experiment was conducted in spring 2014 with cod from the Risør area. Cod were caught, using non-baited fyke nets (mesh size: 20 mm), from late November 2013 to early January 2014 (see Kuparinen, Roney, Oomen, Hutchings, & Olsen, 2016). Individuals larger than 65 cm and smaller than 40 cm were excluded to 1) ensure that the fish were mature, 2) enhance mating prospects, and 3) avoid cannibalism which can arise if size distributions are too broad (Rowe, Hutchings, & Skjæraasen, 2007). This left a total of 73 potentially mature cod for the experiment: 21 females and 16 males from inner Risør and 24 females and 12 males from outer Risør (Figure 1). Cod were individually marked upon capture, using T-bar anchor tags (Hallprint Pty. Ltd., South Australia) and a standard tag applicator. Immediately after capture the cod were transported to the *Institute of Marine Research, Flødevigen*, approximately 60 km from Risør. Adults were held together in a 45 m^3^ spawning basin lined with natural rock and allowed to spawn undisturbed. Daily observations of egg production (from a collection box located at the surface outflow of the basin) showed that the fish started spawning in February and continued spawning until mid-April. The spawning basin and the surrounding environment reflected ambient temperature and photoperiod (for details see Roney et al., 2018).

### Experimental design

Eggs for the experiment were collected at the peak of spawning (i.e., day 45 out of a 94-day spawning season, on which day the greatest number of eggs were observed; Figure S1), to maximize the number and diversity of families while minimizing variation in time since fertilization. The eggs were collected and held in a 900 L flow-through seawater incubation tank at 6°C until hatch, when they were haphazardly sampled and transferred to 40 L flow-through seawater experimental tanks. There were 12 tanks in total, each containing 2000 larvae. The larvae were reared at three temperatures with four replicate tanks per temperature. The target temperatures were 6°C, 9.5°C and 13°C, representing: 1) the average temperature outside the research facility in March/April (i.e., during and immediately after peak spawning); 2) a 3.5°C increase consistent with the projected increase of 2-4°C by the year 2100 (IPCC, 2013) or the average temperature outside the research facility in April/May (i.e., ecologically relevant temperatures for offspring of late spawners); and 3) a 7°C increase consistent with the projected 2-4°C increase relative to the average temperature outside the research facility in April/May (i.e., projected temperatures for offspring of late spawners) (Table S1). Temperatures were measured daily, with average (±SD) temperatures of 6.3 ± 0.2°C, 9.7 ± 0.2°C, and 13.3 ± 0.3°C in the low, intermediate and high temperature tanks, respectively. Water quality parameters (oxygen, pH, and ammonia concentration) were monitored with no notable deviations. The larvae were fed rotifers (*Brachionus plicatilis*) enriched with RotiGrow *Plus*™ (Reed Mariculture, USA) in excess (4500 prey/L three times daily) (Hutchings et al., 2007). The experiment was terminated at 28 days post hatch (dph), at which time the remaining larvae were counted prior to sampling as a measure of total survival.

### Sample collection

Forty larvae were sampled haphazardly from each tank replicate at 2 and 28 dph, as well as forty larvae from the incubation tank at 0 dph (n=1000). Forty larvae were also removed from each tank at 14 and 21 dph for purposes outside of the present study (n=960). Samples collected at 0 and 2 dph were used to assess initial lengths. Length at 28 dph is used as a proxy for growth, following Hutchings et al. (2007). Sampled larvae were put on a glass slide containing RNAlater (Ambion), immediately photographed using a stereoscope and a Leica DFC 425 C camera, and submerged in a 1.5 ml microtube containing 250 μl RNAlater. The photographs were used to measure standard larval length (from the tip of the longest jaw to the end of the notochord; Kahn et al., 2004), using the segmented line tool in the software ImageJ (Abràmoff, Magalhães, & Ram, 2004). A 1 mm scale bar photographed at the same time as the larvae was used for calibration. Three larvae were excluded because of missing or poor-quality photographs. Some larvae were unable to be photographed in a straight position within the first few seconds of RNAlater exposure. Therefore, larval curvature was defined as 0=not curved, 1=slightly curved, and 2=very curved in order to statistically account for potential measurement bias of curved individuals. A subset of larvae (n=120) was measured twice and the correlation coefficient was used to infer that measurement error was low (slope = 1.0054, R^2^ = 0.99).

### DNA isolation and microsatellite genotyping

Of the 1000 larvae sampled, 874 were selected for genotyping from 3 (28 dph) or 4 (2 dph) tank replicates. DNA was individually extracted from whole larvae, using the E-Z 96 DNA/RNA Isolation Kit (Omega Bio-Tek, USA), and from parental fin clips, using E.Z.N.A. Tissue DNA Kit according to the manufacturer’s protocol for tissue samples with elution buffer preheating. One well per plate was used as a negative control.

PCR amplification was performed in two multiplexes containing four loci each (Table S2). The multiplexes were modified from Delghandi et.al. (2003) (Multiplex 1) and Dahle et al. (2006) and Glover et al. (2010) (Multiplex 2), and were previously used for parental assignment of offspring from the same broodstock as that used in the present study (Roney, Oomen, Knutsen, Olsen, & Hutchings, 2018b).

The first multiplex consisted of 1.50 mM buffer, 0.30 mM dNTP, 0.80 U QiagenTaq pol, 0.12 µM GMO19, 0.32 µM TCH11, 0.04 µM GMO8, 0.20 µM GMO35 (all primers one forward and one reverse) and distilled H2O. The second multiplex consisted of 1.50 mM buffer, 0.3 mM dNTP, 0.80 U QiagenTaq pol, 0.12 µM GMO34, 0.23 µM GMO132, 0.18 µM GMO2, 0.35 µM TCH13 (forward and reverse) and distilled H2O. The total volume of each PCR was 10 µl, of which 1 µl was DNA extract of unknown concentration.

The PCR cycling conditions for both multiplexes consisted of an initial 5 min denaturation at 95°C followed by 30 cycles of denaturation (30 s at 90°), annealing (90 s at 56°C), and extension (60 s at 72°C), and a final 10 min extension step. PCR products were held at 4°C before being visualized using an ABI PRISM 3130xl Genetic Analyzer (Applied Biosystems). Genotypes were scored using Genemapper software v.4.0 (Life Technologies). To ensure accuracy of parental genotypes, all adults were amplified three times (for details see Roney et al., 2018b), as were individuals with 3 or more missing loci or ambiguous genotypes. Individuals were independently genotyped by two (larvae) or three (adults) researchers. In Genemapper, default analysis settings were applied (minimum peak height 50 and size standard GS500LIZ). All adults and 868 larvae were successfully genotyped at 4 or more microsatellite loci.

### Parentage analyses

Heterozygosities at each locus (H_o_=observed heterozygosity within samples, H_e_=expected heterozygosity total loci (Nei & Chesser, 1983)) were estimated for the adults, using Genetic Data Analysis software (Lewis & Zaykin, 2001) (Table S2). F_IS_-values were estimated for the adults in GENEPOP v.4.2 (Rousset, 2008). Tests for deviations from Hardy-Weinberg equilibrium (Weir & Cockerham, 1984), conducted using the exact probability test with false discovery rate (FDR; α=0.05) correction for multiple testing (Benjamini & Hochberg, 1995), showed minimal deviations (Table S3). These statistics were performed on adult genotypes only. CERVUS v3.0 (Kalinowski, Taper, & Marshall, 2007) was used to assign parents to offspring, using allele frequencies based on the adults. To estimate genotyping error, 78 larvae were amplified twice and the genotypes were compared, resulting in an error rate of 0.006536. For a subset of individuals from 2 and 28 dph, assignment was run with both the estimated error rate and a global error rate of 0.01 and the resulting assignments were identical. Therefore, a global error rate of 0.01 was used to analyze the complete data set. The proportion of loci typed was 0.9943, and the minimum number of typed loci was four. For all other parameters, default settings were used. Parental assignments were made based on two criteria: 1) a confidence level of >95%, estimated from a simulated analysis of 10,000 offspring, and 2) a maximum of 1 mismatched locus per trio or two mismatched loci if both were only one repeat away from a putative parent allele (n=11). The second criterion was implemented because CERVUS forces an assignment if all parent genotypes are known.

Parental assignment identified 10 mothers (6 inner and 4 outer fjord) and 14 fathers (10 inner and 4 outer fjord) (Table S4). Larvae were classified into crosses depending on whether their mother and father were collected from inner (I) or outer (O) Risør: (mother×father) = I×I, I×O, O×I, and O×O. These crosses consisted of 236, 257, 322, and 51 larvae belonging to 12, 7, 8, and 5 families, respectively (Table S5).

### Parental origin assignments

To discriminate the parental ecotype (fjord or North Sea), all adults were genotyped with a Sequenom MassARRAY multiplex of 40 single nucleotide polymorphisms (SNPs) specially developed to distinguish between these ecotypes (Jorde, Kleiven, et al., 2018; Knutsen et al., 2018). The SNP panel was developed based on reference samples from 1) mature adult cod in the North Sea (n=91), and 2) juvenile (0+) cod from the innermost areas of three Skagerrak fjords, including Risør (n=143) (for details see Jorde, Kleiven, et al., 2018). Briefly, 9,187 SNPs from a 12k SNP array were scored in the reference samples as well as in cod caught by commercial trawls outside of the aforementioned Skagerrak fjords (n=118) (Jorde, Kleiven, et al., 2018; Sodeland et al., 2016). SNPs were ranked by Nei’s (1973) GST between fjord and North Sea samples and filtered to exclude SNPs that were highly linked. A composite linkage disequilibrium (CLD; Gao et al., 2008) >0.5 to a higher ranked SNP resulted in exclusion. After filtering, 40 high-*G*ST-ranked SNPs were selected for genotyping in the multiplex. In our study and that of Knutsen et al. (2018), 26 of these SNPs were scored successfully. Assignments of the adult cod in our experiment to the two reference samples were performed with Geneclass II v.2.0 (Piry et al., 2004).

Individuals with fewer than 20 scored SNPs (n=2) were excluded from further consideration. Assignments with a score <80% (n=3) were considered to be ambiguous and the remaining assignments scored >90%. Some of the offspring of fertilizations between one individual with ambiguous assignment (RIC5100) and unambiguously fjord type cod was later analyzed using a larger set of SNPs and found to cluster with fjord × North Sea hybrid offspring (Figure S2). Therefore, RIC5100 was classified as a North Sea ecotype. The remaining individuals with ambiguous assignment did not produce offspring contributing to this experiment. In sum, ecotype assignments were obtained for 69 of 73 adults present in the spawning basin, including all of those identified as parents in the present study (Table S4).

### Growth reaction norms

Variation in plasticity is reflected in non-parallel reaction norms, which are graphical representations of the range of phenotypes expressed by a genotype along an environmental gradient (Woltereck 1909; Schmalhausen 1949). All statistical analysis on growth and survival reaction norms were conducted in R v.3.2.3 (R Development Core team 2015). A linear mixed effects model implemented with the lme4 package (Bates, Mächler, Bolker, & Walker, 2015) was used to test for differences in length between crosses at the beginning of the experiment (0 and 2 dph combined), with cross as a fixed effect, larval curvature as a covariate, and the individual identity of the mother and father as random (i.e., parental) effects [1].

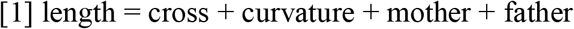

Differences in thermal growth reaction norms between crosses were tested using a mixed effects linear model on length at 28 dph, with cross, temperature, and their interaction as fixed effects, larval curvature as a covariate, ecotype and tank as random effects nested within temperature, and mother and father as random effects [2]. Temperature was modelled as a factor to allow for non-linear responses.

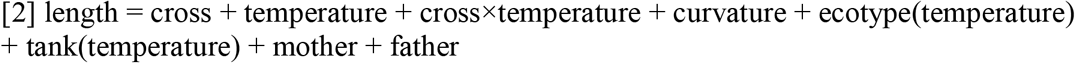

A significant cross×temperature interaction indicates a significant difference in the slopes of the reaction norms and suggests that there are genetic differences in thermal responses between crosses (i.e., genetic variation in plasticity). Post-hoc contrasts were used to test for 1) temperature effects (i.e., plasticity) within crosses and 2) differences in slopes (i.e., variation in plasticity) between crosses. Contrasts were examined for the lower (6-9.5°C) and upper (9.5-13°C) range of temperatures separately. Length at 28 dph followed a normal distribution (Figure S3) and variances were fairly homogeneous among treatments, as assessed by visual inspection of the model residuals (Figure S4, S5).

To assess the robustness of our growth model to ecotype, we repeated the analysis without ecotype as a random effect and obtained similar results (Table S6; Figure S6).

### Survival reaction norms

Cross-specific survival was quantified for each tank at the end of the experiment (28 dph). It was not feasible to genotype all surviving individuals, but rather only those that were sampled for growth measurements. Therefore, survival was estimated for each cross and tank using the following formula, where 40 is the total number of larvae sampled for genotyping from each tank, n_c_ is the number of genotyped larvae assigned to each cross from each sample, and n_t_ is the total number of larvae alive in the tank at the end of the experiment:

Survival = [n_c_/40]*n_t_

Values were rounded to the nearest integer. We did not correct for sampling mortality ([n_2dph_+n_14dph_+n_21 dph_]/initial n = [40+40+40]/2000 = 6%), as the same number of larvae were removed from each tank.

We tested for differences in survival reaction norms between crosses, using a generalized linear model with a quasi-poisson distribution (dispersion parameter = 15.3 >1) and identity link. Cross, temperature, and their interaction were included as fixed effects [3].

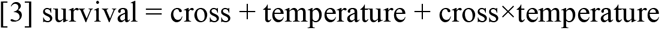

Deviances were evaluated with Chi square tests and the forward stepwise method. Post-hoc contrasts were used to identify the differences. The model residuals were normally distributed, as assessed by visual inspection (Figure S7). The variance was slightly lower at 13°C but all variances were evenly distributed around zero (Figure S8, S9).

## Results

### Genetic assignments

Adult cod collected from inner Risør were dominated by the fjord ecotype (91.2%) and outer Risør cod were dominated by the North Sea ecotype (80.0%). Of those identified as parents in our study, the North Sea ecotype was only detected in outer Risør individuals. The majority of offspring (66%; n=570) were pure Fjord ecotype. Offspring representing hybrids between fjord and North Sea ecotypes were also common (34%; n=295), whereas only one offspring (0.1%) was pure North Sea ecotype.

### Growth reaction norms

There was no effect of cross on initial length (*P*=0.831; Table 1a), for which mother and father explained 3% and 0% of the variance, respectively. However, there was a significant cross×temperature interaction for length at 28 dph (*P*=0.005; Table 1b), indicative of variation in thermal plasticity for growth between crosses (Figure 2). Growth generally increased from 6°C to 9.5°C, although this effect was only significant for I×I and I×O (Table 2). However, from 9.5°C to 13°C, growth decreased for all crosses except for O×I (Table 2). Again, this effect was only marginally significant for I×I and I×O yet produced significant variation in reaction norm slopes between these crosses and O×I across the warmer temperature range (Table 3). O×O also had a marginally different slope compared to O×I across warmer temperatures (Table 3). As a result, minimal variation in mean length was observed between crosses at the coldest temperature, whereas up to a 1.1 mm (15%) difference in mean length was observed at the warmest temperature (Figure 2). This difference at the warmest temperature corresponded to a 4-12% decrease in size (relative to the maximum) for I×I, I×O, and O×O, but a 4% increase in size for O×I. Mother and father explained 1% and 3% of the variance in growth, respectively, whereas ecotype explained 7%, 3%, and 10% at 6°C, 9.5°C, and 13°C, respectively (Table 1b).

**Table 1:** Results of linear mixed effects models for (a) initial larval length (0 and 2 dph) and (b) growth (28 dph) for 4 crosses of Atlantic cod. Asterisks denote significance at α=0.05.

| (a) Initial length |  |  |  |  |  |
| --- | --- | --- | --- | --- | --- |
| Model term | d.f. | Sum of squares | Mean of squares | F | P-value |
| cross | 3 | 0.06 | 0.02 | 0.47 | 0.831 |
| curvature | 1 | 0.10 | 0.10 | 2.53 | 0.112 |
| Model term | Variance | SD |  |  |  |
| father | 0.00 | 0.01 |  |  |  |
| mother | 0.03 | 0.17 |  |  |  |
| residual | 0.04 | 0.20 |  |  |  |
| (b) Growth |  |  |  |  |  |
| Model term | d.f. | Sum of squares | Mean of squares | F | P-value |
| cross | 3 | 0.40 | 0.13 | 0.39 | 0.764 |
| temperature | 2 | 0.78 | 0.39 | 1.12 | 0.327 |
| curvature | 1 | 6.07 | 6.07 | 17.48 | <0.001 * |
| cross×temperature | 6 | 6.63 | 1.11 | 3.18 | 0.005 * |
| Model term | Variance | SD |  |  |  |
| father | 0.03 | 0.18 |  |  |  |
| mother | 0.01 | 0.11 |  |  |  |
| ecotype at 6°C | 0.07 | 0.26 |  |  |  |
| ecotype at 9.5°C | 0.03 | 0.16 |  |  |  |
| ecotype at 13°C | 0.10 | 0.31 |  |  |  |
| tank | 0.01 | 0.12 |  |  |  |
| temperature:tank | 0.02 | 0.15 |  |  |  |
| residual | 0.35 | 0.59 |  |  |  |

**Table 2:** Effect of increasing temperature on larval growth for 4 crosses of Atlantic cod. Asterisks denote significance at the following levels of α: *0.10 and **0.05.

| Cross | Estimate | SEM | % | *t* | *P*-value |  |
| --- | --- | --- | --- | --- | --- | --- |
| *6-9.5°C* |  |  |  |  |  |  |
| I×I | 0.45 | 0.26 | 3.97 | 1.74 | 0.046 | ** |
| I×O | 0.52 | 0.27 | 8.12 | 1.95 | 0.031 | ** |
| O×I | 0.26 | 0.25 | 3.12 | 1.06 | 0.148 |  |
| O×O | -0.07 | 0.33 | 2.25 | -0.20 | 0.422 |  |
| *9.5-13°C* |  |  |  |  |  |  |
| I×I | -0.49 | 0.31 | -4.26 | -1.60 | 0.062 | * |
| I×O | -0.47 | 0.32 | -8.67 | -1.47 | 0.071 | * |
| O×I | 0.11 | 0.27 | 4.21 | 0.40 | 0.345 |  |
| I×O | -0.51 | 0.42 | -12.36 | -1.21 | 0.136 |  |

**Table 3:** Pairwise contrasts of the effect of increasing temperature on larval growth for 4 crosses of Atlantic cod. Estimates (±SEM) are given above the diagonal and P-values below. The point of contrast for the estimates is the row header. Asterisks denote significance at the following levels of α: *0.10 and **0.05.

| 6-9.5°C | I×I | I×O | O×I | O×O |
| --- | --- | --- | --- | --- |
| I×I | - | 0.075 (0.257) | -0.187 (0.191) | -0.515 (0.338) |
| I×O | 0.387 | - | -0.261 (0.251) | -0.590 (0.300) |
| O×I | 0.168 | 0.154 | - | -0.329 (0.330) |
| O×O | 0.078* | **0.037\*\*** | 0.170 | - |
| 9.5-13°C | I×I | I×O | O×I | O×O |
| I×I | - | 0.024 (0.320) | 0.601 (0.203) | -0.018 (0.442) |
| I×O | 0.470 | - | 0.576 (0.292) | -0.042 (0.382) |
| O×I | **0.003\*\*** | **0.034\*\*** | - | -0.619 (0.419) |
| O×O | 0.484 | 0.458 | 0.095* | - |

**Figure 2:**
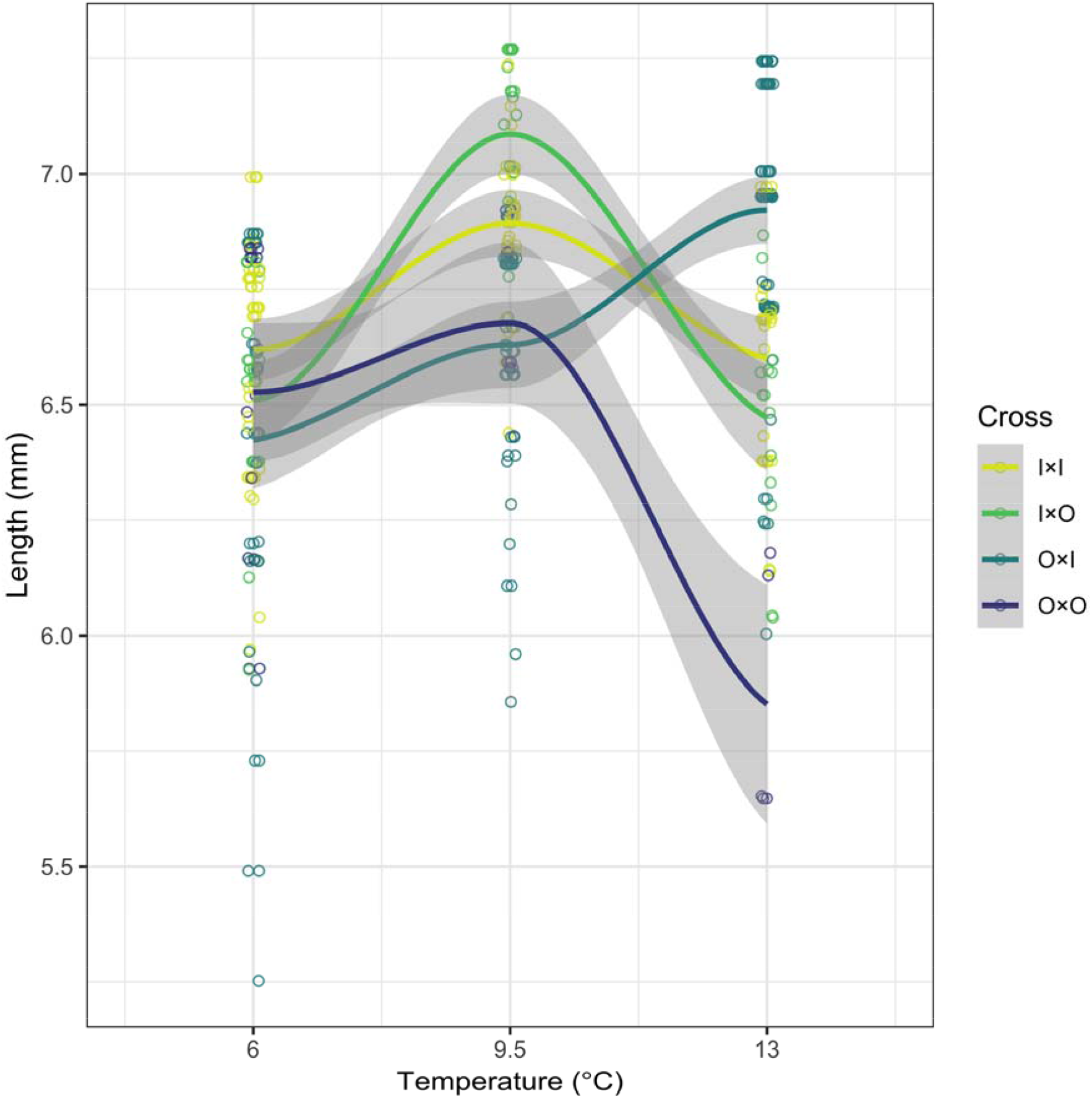
Thermal reaction norms for larval Atlantic cod growth for four crosses (mother×father: I = inner Risør fjord, O = outer Risør fjord) based on a linear mixed effects model with smoothed conditional means calculated from a loess regression on the model-fitted lengths. Shading represents 95% confidence intervals.

### Survival reaction norms

There were significant effects of cross (*P*=0.002), temperature (*P*<0.0001), and a cross×temperature interaction (*P*=0.002) for survival, indicative of variation in thermal plasticity between crosses (Figure 3, Table 4). All crosses lacked significant plasticity from 6°C to 9.5°C, although there was a marginal increase in survival of O×I (Figure 3, Table 5). In contrast, all crosses showed a significant decrease in survival at the highest temperature, with reductions of 76-93% from 9.5°C to 13°C (Figure 3, Table 5). There were significant differences in the magnitudes of these reductions, with I×I, I×O, and O×I exhibiting steeper declines in survival at the highest temperature compared to O×O (Table 6). The rank survival was generally O×I, I×I, I×O, and O×O from highest to lowest, except for O×I and I×I which had similar survival at the coldest temperature (Figure 3).

**Table 4:** Deviance table from generalized linear models for the effects of cross and temperature on larval Atlantic cod survival. The P-values were obtained from χ2 tests of whether the model fit improved by sequentially adding population, temperature, and their interaction to the null model. Asterisks denote significance at α=0.05.

| Model term | d.f. | Deviance | Residual d.f. | Residual deviance | *P*-value |  |
| --- | --- | --- | --- | --- | --- | --- |
| null |  |  | 35 | 2329.30 |  |  |
| cross | 3 | 685.66 | 32 | 1643.60 | 0.002 | * |
| temperature | 2 | 978.66 | 30 | 664.96 | <0.001 | * |
| cross×temperature | 6 | 312.14 | 24 | 352.82 | 0.002 | * |

**Table 5:** Effect of increasing temperature on larval survival for 4 crosses of Atlantic cod. Asterisks denote significance at the following levels of α: *0.10 and **0.05.

| Cross | Estimate | SEM | % | *t* | *P*-value |  |
| --- | --- | --- | --- | --- | --- | --- |
| *6-9.5°C* |  |  |  |  |  |  |
| I×I | 26.05 | 38.87 | 16.22 | 0.67 | 0.509 |  |
| I×O | 30.55 | 32.81 | 25.37 | 0.93 | 0.361 |  |
| O×I | 76.54 | 42.07 | 36.25 | 1.82 | 0.081 | * |
| O×O | 4.52 | 20.79 | 10.16 | 0.22 | 0.830 |  |
| *9.5-13°C* |  |  |  |  |  |  |
| I×I | -145.79 | 29.97 | -90.77 | -4.87 | <0.001 | ** |
| I×O | -112.50 | 25.64 | -93.40 | -4.39 | <0.001 | ** |
| O×I | -160.45 | 36.61 | -75.99 | -4.38 | <0.001 | ** |
| O×O | -40.26 | 15.78 | -90.57 | -2.55 | 0.018 | ** |

**Table 6:**
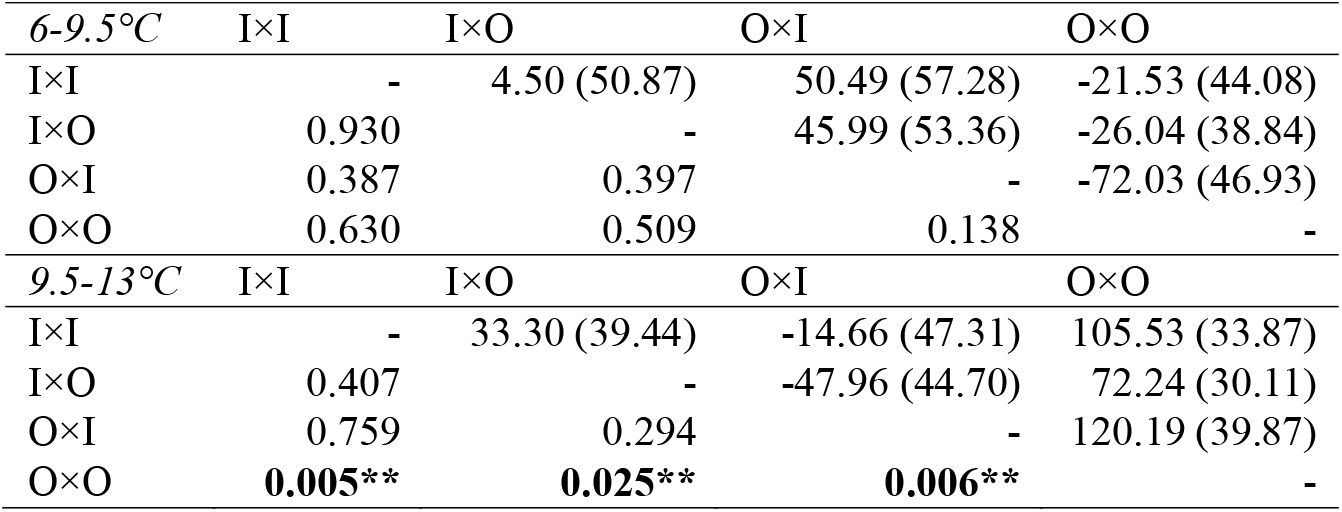
Pairwise contrasts for the effect of increasing temperature on larval survival for 4 crosses of Atlantic cod. Estimates (±SEM) are given above the diagonal and P-values below. The point of contrast for the estimates is the row header. Asterisks denote significance at the following levels of α: *0.10 and **0.05.

**Figure 3:**
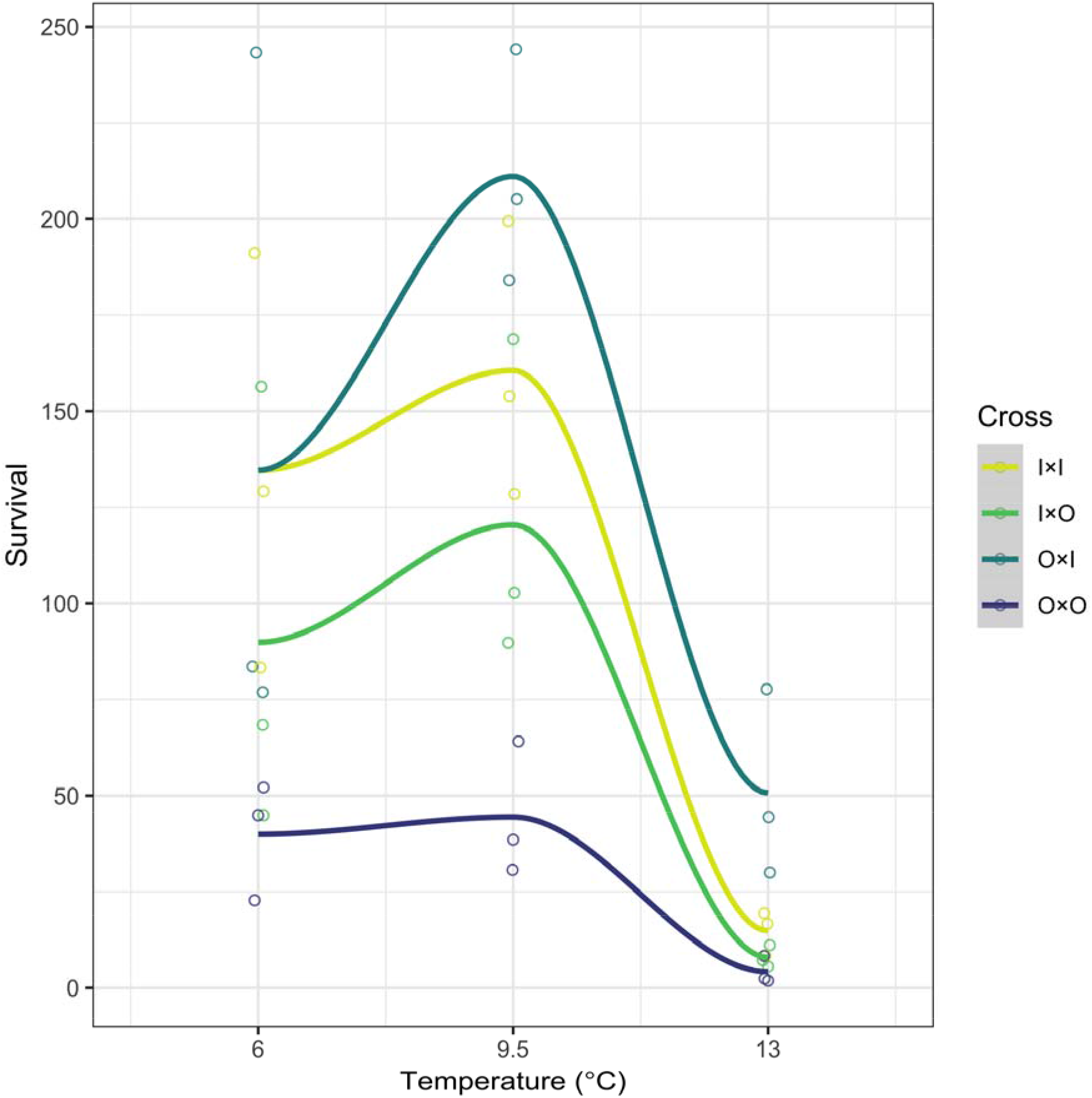
Thermal reaction norms for larval Atlantic cod survival for four crosses (mother×father: I = inner Risør fjord, O = outer Risør fjord) based on a generalized linear model with a quasibinomial distribution. Model-fitted values for each tank replicate (n=3) are shown.

## Discussion

### Divergence in growth and survival plasticity

Here, we shed light on genetically based phenotypic variation of Atlantic cod within a fjord system. We show genetic variation in thermal plasticity for larval growth and survival (i.e., variation in reaction norm slopes), wherein divergence in thermal responses is greater at warmer temperatures. We previously showed that temperatures of 9°C to 13°C elicit a stress response in cod larvae originating from nearby Flødevigen, Norway when reared under similar conditions (Oomen et al., in review), supporting the possibility that these temperatures could be stressful for Skagerrak cod larvae. In the present study, all crosses experienced severe (75-93%) declines in survival and variable growth responses across the same temperatures. This finding suggests that, although slower growth can confer energy savings, which could be beneficial under stressful conditions (Billerbeck, Lankford, & Conover, 2001; Conover & Present, 1990), any potential energy savings from the slower growth of some crosses had a minimal impact on fitness. However, the cross with the greatest reduction in growth at 13°C experienced the smallest reduction in survival at warm temperatures (O×O; Table 2, Table 5), which is consistent with energy savings benefits. Plasticity that results in smaller sizes might also be maladaptive, as this can extend the duration of the vulnerable larval phase (Anderson, 1988).

While lower temperatures currently characterize the bulk of the spawning and subsequent larval seasons (Table S1), temperatures can increase rapidly in late spring, with high year-to-year variability (Olsen, Carlson, Gjøsæter, & Stenseth, 2009; Oomen et al., in review). In May 2014, shortly after termination of the present study, temperatures in the Risør fjord area averaged 10-12°C with maximums of 12-16°C at 5 m and 10 m depths (Table S1). Therefore, it is probable for the offspring of late (i.e., late April-May) spawners to encounter warm enough temperatures to experience severe reductions in survival. Further, an increase in sea temperature of 2-4°C, as projected to occur before the year 2100 (IPCC, 2013), would increase exposure of the offspring of peak-season (i.e. March-early April) spawners to such warm temperatures. The decreased larval fitness associated with warmer temperatures documented in the present study might reduce population productivity and connectivity of Skagerrak coastal cod.

High larval mortality in cod is common in the wild (Houde & Zastrow, 1993; Sundby, Bjørke, Soldai, & Olsen, 1989) and in the laboratory (Gamble & Houde, 1984; Oomen & Hutchings, 2015b, 2016; Otterlei, Nyhammer, Folkvord, & Stefansson, 1999; Steinarsson & Björnsson, 1999). If size-selective mortality was responsible for the variation in growth plasticity observed, we would expect to see a consistent correlation between changes in length and survival. However, the cross with the smallest reduction in survival from 9.5°C to 13°C (O×O) had a similar growth slope to some (I×I and I×O), but not all (O×I), crosses that experienced higher mortality (Table 3). Therefore, variation in growth is not explained by variation in survival in our experiment.

### Evolutionary potential in a warming ocean

Greater divergence in reaction norms at the highest temperature is consistent with the hypothesis that extreme environments reveal cryptic genetic variation (Ghalambor et al., 2007; Murren et al., 2014). Importantly, significant genetic variation in survival plasticity suggests that some larvae are more vulnerable to mortality at higher temperatures than others and that this trait has a genetic basis. The availability of such standing genetic variation for selection to act upon is key to producing an adaptive response to ocean warming (Bernatchez, 2016), provided that the necessary variants exist in the population and that evolutionary change can keep pace with the environment to avoid extinction (Maynard Smith, 1989; Merilä & Hendry, 2014). In this way, natural genetic variation across heterogeneous environments can facilitate rapid evolutionary responses to environmental disturbance (Therkildsen et al., 2019).

The severe and ubiquitous increase in mortality we observed at 13°C is likely to reduce population viability under ocean warming. The probability of persistence will depend on the abundance of well-suited genotypes (Orr & Unckless, 2008) and the potential for gene flow of beneficial alleles into local coastal areas that could effect a ‘genetic rescue’ (Tallmon, Luikart, & Waples, 2004). Coastal cod in Norway is at or near historically low levels (ICES, 2018), reducing the copy number of potentially beneficial alleles. Further, gene flow is reduced among coastal areas due to spawning site fidelity (Espeland et al., 2007), natal homing (André et al., 2016; H. Svedäng, Righton, & Jonsson, 2007), sedentary behaviour (Espeland et al., 2008; Rogers et al., 2014; Villegas-Ríos, Réale, Freitas, Moland, & Olsen, 2017, 2018), and the retention of pelagic larval stages inside fjords (Ciannelli et al., 2010), which could reduce the likelihood of genetic rescue.

While there is a large influx of larvae from the North Sea to coastal Skagerrak (Knutsen et al., 2004; Stenseth et al., 2006), the maintenance of divergent fjord and North Sea ecotypes suggests reduced gene flow between them and reduced fitness of the North Sea ecotype in the inner fjord environment (Knutsen et al., 2018), as supported by survival analysis of tagged fish (Barth et al., 2019). Further, Skagerrak coastal cod are already near the upper thermal limit of the species distribution (Björnsson & Steinarsson, 2002; Righton et al., 2010). Therefore, it is less likely that nearby populations will harbour unique, locally adapted alleles conferring tolerance to higher temperatures. Conversely, the flow of alleles that are well suited to warm temperatures but poorly suited to the local coastal environments could lead to outbreeding depression (Tallmon et al., 2004). Under these conditions, it will be important for local populations to possess sufficient abundances of adaptive alleles in order to adapt to rising water temperatures, through a greater abundance of individuals and conservation of genetic diversity (Hoffmann, Sgrò, & Kristensen, 2017).

### The coastal zone is a potential hybrid zone

The ecotype proportions among the adult cod collected from inner and outer Risør fjord were consistent with a long-term survey of juvenile ecotype composition in this fjord system (Knutsen et al., 2018). We show that, provided the opportunity in semi-natural laboratory conditions, there is potential for substantial reproduction between cod inhabiting the inner and outer parts of Risør fjord (previously shown by Roney, Oomen, Knutsen, Olsen, & Hutchings, 2018b) and between cod of fjord and North Sea ecotype (first documented here). The range of the fjord ecotype is nearly completely overlapped by that of the North Sea ecotype in coastal areas and the presence of eggs, larvae, and adults in spawning condition has previously suggested that the North Sea cod reproduce there (Jorde, Kleiven, et al., 2018; Jorde, Synnes, et al., 2018; Knutsen et al., 2018; Rogers et al., 2014). Yet, it remained unclear whether these ecotypes reproduce with each other, given their temporally stable genetic differentiation (Knutsen et al., 2018). We provide the first direct evidence of the potential for hybridization between ecotypes, including viability of hybrid offspring, thereby providing support against several pre-zygotic isolating mechanisms (temporal, behavioural, and mechanical) and a post-zygotic barrier (hybrid inviability) to gene flow in this system (Rundle & Nosil, 2005).

### Genetic basis of variation in thermal plasticity

Although the focus of the present study was on variation among crosses based on location, it is notable that ecotype explained a minor (3-10%) portion of the variance in growth reaction norms. Therefore, while there is support for a relatively small role of ecotype in shaping thermal growth responses, it is an insufficient descriptor of biological variation in the present study. The precise nature of ecotypic variation could have been explored using a full factorial breeding design instead of semi-natural spawning conditions. However, such a design requires stripping gametes to make artificial crosses, which stresses the parents and therefore can negatively influence offspring fitness. We avoided imposing such an artificial selection pressure on the phenotypes of interest. Further, by reflecting the phenotypic diversity present in the wild, our findings are more evolutionarily relevant for conservation and management.

Nonetheless, the observed variation in thermal plasticity in common environments suggests that genetic variants not examined in the present study affect early life traits and thermal plasticity in coastal cod. One alternative hypothesis is that chromosomal inversions (Sodeland et al., 2016; Berg et al., 2017) are responsible for the variation in thermal plasticity we observed. The inversions are not alternately fixed in fjord and North Sea ecotypes and their specific functions and key environmental drivers remain unclear (Barth et al., 2019). Five out of the twenty-six SNPs used to distinguish between Fjord and North Sea ecotypes were located inside inversions (Knutsen et al., 2018; Sodeland et al., 2016) and their lack of alternate fixation may have contributed to the initially ambiguous population assignments of some of the parental fish in the present study. We do not know the inversion genotypes of the individuals in the present study. Further research is needed to determine whether these inversions play a role in thermal adaptation and plasticity in Atlantic cod.

Epigenetic effects represent another potential source of variation in thermal plasticity, as documented in several fishes (reviewed by Donelson et al., 2018). We found no differences in initial length among crosses, which suggests a lack of variation in maternal effects on egg size among crosses (Marshall, 2008). Although the parents were held for 2-3 months in a common environment, epigenetic effects could contribute to differences between locations. However, there is evidence to the contrary in the form of a lack of correspondence between crosses with either a mother (O×I and O×O) or father (I×I and O×I) from the same location. Because maternal and epigenetic effects do not explain the patterns of variation in thermal responses observed in the present study, as was also the case in previous studies of larval cod plasticity (Oomen & Hutchings, 2015b, 2016), we interpret the variation to be largely of genetic origin. However, the potential contribution of epigenetic effects warrants further investigation.

### Spatial scale of genetic variation

The ∼20 km^2^ Risør fjord represents the smallest spatial scale at which genetic variation in adaptive traits has been examined in Atlantic cod, at an order of magnitude smaller than previously identified adaptive differentiation (Oomen & Hutchings, 2015b). In contrast to our predictions based on contrasting thermal environments in the inner and outer parts of the fjord system, we find greater variation in plasticity within the system as a whole than between inner and outer fjord locations. This is because pure crosses tended to exhibit more similar growth reaction norms to each other than to hybrid crosses. Also contrary to our predictions was a lack of intermediate responses in the hybrids that would be indicative of additive genetic effects. Rather, our observations of divergent, non-additive reaction norms of reciprocal hybrids (e.g., I×O across lower temperatures and O×I across higher temperatures for growth plasticity and O×I for mean survival at both warmer temperatures) provide limited evidence for a hypothesis of hybrid overdominance. Given that there is at least some gene flow between locations in the wild (Barth et al., 2019; Knutsen et al., 2011), this again suggests that there is likely greater variation in thermal responses within the fjord system than between inner and outer locations.

These findings do not preclude local adaptation to thermal regimes in this system, particularly as survival plasticity differed between the pure cross O×O and the remaining crosses that had some genetic component from the inner fjord. The present study contributes to a growing body of work suggesting the presence of adaptive variation within fjord systems in Atlantic cod (Barth et al., 2017; Jorde, Kleiven, et al., 2018; Knutsen et al., 2018; Kuparinen et al., 2016; Ono et al., 2019; Roney, Oomen, Knutsen, Olsen, & Hutchings, 2018a; Sodeland et al., 2016), and is the first to examine the influences of spatial environmental heterogeneity (location) while accounting for ecotype.

We also provide rare evidence of genetic variation in a fitness-related trait in Atlantic cod. Knutsen et al. (2018) found differences in juvenile growth rates between ecotypes within similar habitats in the wild, although the genetic basis of this is unconfirmed. Using the same parental fish as the present study, Kuparinen et al. (2016) determined that cod from outer Risør had higher adult growth rates than cod from inner Risør. However, the extent to which differences in ecotype frequencies between locations might have contributed to this trend, and the relative contributions of genotype and environment to growth in the wild, are unknown. A related study demonstrated higher reproductive success in the cod from inner Risør compared to outer Risør, after accounting for differences in size and age (Roney et al., 2018a). Although ecotype was not explicitly considered, this finding suggests a potential reproductive barrier between cod from inner and outer Risør. Such a barrier could promote adaptive divergence within fjords, especially when combined with environmental differences between fjord and outer coastal areas. Yet altogether, the existing body of work suggests that local adaptation in this system is likely complex and nuanced, with differentiation across multiple axes of genomic variation (e.g., neutral and adaptive SNPs, structural variants) and novel combinations arising from mixing between spawning components.

### Conservation and management of mixed-ecotype coastal fisheries

While the present study provides limited evidence of local adaptation, it highlights the presence of functional genetic variation in Skagerrak coastal cod beyond ecotype and the need for a better understanding of the genomic basis of the variable thermal responses observed. We further raise the potential for extreme temperatures and hybridization to generate diversity in thermal responses for selection to act upon, which underscores the need for a management strategy that protects the intraspecific genetic diversity necessary for maintaining resilient local populations. When the genetic components of a population differ in abundance or adaptive traits, including responses to environmental variables, strategies that fail to account for this diversity are at a higher risk of failure (Schindler et al., 2010). Considering that many marine fishes are overexploited (Costello et al., 2012; Worm et al., 2009) and that ocean warming is occurring rapidly (IPCC, 2013), resolving the spatial genetic substructure of harvested fishes is of considerable importance.

Regardless of the specific genetic basis of adaptive variation, the ubiquitous negative impact of 13°C or less on larval cod fitness and its likely impact on recruitment suggest that regulations will need to be adjusted as the sea warms. Currently, commercial fisheries within 12 nautical miles of the Norwegian Skagerrak coast are only subject to technical regulations (e.g., minimum landing size, mesh size), not catch limits, like those in place just outside of the coastal zone (Jorde, Kleiven, et al., 2018). These regulations should be re-examined in the context of projected warming scenarios and rapid-response measures should be planned in case of an abrupt increase in temperature like that observed in the late 1980s (Olsen et al., 2009). Likewise, management plans, which are currently updated on decadal timescales (ICES, 2018), will probably need to be revised at a finer temporal scale given the rapid pace of global warming.

The high, yet diverse, thermal sensitivities of cod larvae underscore the need for enhanced monitoring of local populations. Although recent decades have seen declines in populations along the Skagerrak coast (Olsen et al., 2009; Roney et al., 2016; Svedäng & Bardon, 2003), relatively little is known about the productivity of the fjord components, as they have previously been assumed to be identical to those occupying the outer coastal areas. The fjord ecotype is more likely to be targeted by the recreational fishery (for which neither catch nor effort data are collected) in this densely populated summer destination (Jorde, Kleiven, et al., 2018; Kleiven et al., 2016). Poor documentation of both recreational and commercial cod fisheries along the coast (Jorde, Kleiven, et al., 2018; Kleiven et al., 2016) fundamentally impedes the ability of fisheries management to respond in a changing environment. Improvements in monitoring of both fishing activities and local population productivities represent, one would think, relatively agreeable actions for all stakeholders. A recent ban on fishing cod along much of the Skagerrak coast underscores the urgency of need for data to inform a sustainable management strategy. In general, broad-scale management and lack of coastal monitoring impede the development of strategies to maintain the potential to adapt to ocean warming in systems with phenotypic complexity resulting from cryptic genetic variation, coexisting ecotypes, and gene flow.

## Data Archiving Statement

Data for this study are available at: to be completed after manuscript is accepted for publication.

## Acknowledgements

We acknowledge the contributions of the staff and students at the *Institute of Marine Research, Flødevigen* for their facilities, technical assistance, and additional support during the experiment, especially P. Baardsen, S. Stiansen, R. Johansen, H. Sannaes, and K. Enersen. Special thanks to N. Roney and D. Rivas-Sánchez for help with the experiment and to A. Slettan for lending us the MagNA Lyser. We are grateful to the Centre for Ecological and Evolutionary Synthesis (CEES) for technical and administrative assistance. Thanks to P. Bentzen, C. Herbinger, A. Vøllestad, and L. Bernatchez for helpful comments and discussion. The work was supported by funding from the Natural Sciences and Engineering Research Council of Canada (Discovery Grant to JAH and Canada Graduate Scholarship to RAO), the Norwegian Research Council (HAVKYST Grant to JAH and Leiv Eiriksson Mobility Grant to RAO), Interreg IV (MarGen) to HK, the Ministry of Trade, Industry and Fisheries, Norway, and a Killam Predoctoral Scholarship, Nova Scotia Graduate Scholarship, and James S. McDonnell Foundation Postdoctoral Research Fellowship to RAO.

## Notes

### Competing Interest Statement

The authors have declared no competing interest.

