## Supplementary information for "Cryptic microgeographic variation in responses of larval Atlantic cod to warmer temperatures"

### Supplementary Materials

#### Supplementary Tables

Table S1: Temperatures (°C) in inner and outer Risør fjord at 5 and 10 m depths from January to May 2014.

| | Inner Risør | | Outer Risør | |
|---|---|---|---|---|
| Month | 5 m | 10 m | 5 m | 10 m |
| *Mean* | | | | |
| January | 6.27 | 7.38 | 2.33 | 2.22 |
| February | 2.96 | 4.09 | 2.78 | 2.95 |
| March | 4.48 | 5.69 | 4.82 | 4.86 |
| April | 7.22 | 6.95 | 7.39 | 6.67 |
| May | 12.04 | 9.93 | 11.57 | 10.12 |
| *Standard deviation* | | | | |
| January | 0.08 | 0.08 | 0.08 | 0.06 |
| February | 1.31 | 1.93 | 0.84 | 0.84 |
| March | 1.57 | 1.41 | 0.68 | 0.65 |
| April | 1.65 | 0.81 | 1.48 | 0.87 |
| May | 1.54 | 1.62 | 2.10 | 2.32 |
| *Minimum* | | | | |
| January | 6.06 | 7.28 | 2.09 | 2.09 |
| February | 1.76 | 1.98 | 1.76 | 1.66 |
| March | 2.30 | 2.94 | 3.68 | 3.79 |
| April | 5.76 | 5.86 | 5.66 | 5.55 |
| May | 7.48 | 7.08 | 7.28 | 6.88 |
| *Maximum* | | | | |
| January | 6.37 | 7.48 | 2.41 | 2.30 |
| February | 6.37 | 7.38 | 5.04 | 4.93 |
| March | 7.78 | 7.88 | 6.06 | 5.86 |
| April | 11.53 | 9.47 | 11.63 | 9.08 |
| May | 15.95 | 11.72 | 16.43 | 15.09 |

Table S2. Summary statistics of microsatellite loci used for parentage analysis of Risør cod: number of alleles ($N_{(a)}$), size range (bp), expected heterozygosity ($H_e$), observed heterozygosity ($H_o$), and inbreeding coefficient ($F_{IS}$).

| Locus | N (a) | Size (bp) | Reference | He (Outer) | Ho (Outer) | $F_{IS}$ (Outer) | He (Inner) | Ho (Inner) | $F_{IS}$ (Inner) |
|---|---|---|---|---|---|---|---|---|---|
| GMO8[1] | 23 | 110-205 | Miller et al. 2000 | 0.953 | 1.000 | -0.050 | 0.881 | 0.865 | 0.018 |
| GMO19[1] | 24 | 120-220 | Miller et al. 2000 | 0.930 | 0.833 | 0.106 | 0.919 | 0.784 | 0.149 |
| GMO35[1] | 12 | 110-145 | Miller et al. 2000 | 0.826 | 0.750 | 0.093 | 0.827 | 0.865 | -0.046 |
| TCH11[1] | 20 | 121-193 | O'Reilly et al. 2000 | 0.917 | 0.806 | 0.123 | 0.937 | 0.946 | -0.009 |
| GMO2[2] | 14 | 102-138 | Brooker et al. 1994 | 0.887 | 0.833 | 0.061 | 0.838 | 0.784 | 0.065 |
| GMO34[2] | 8 | 80-120 | Miller et al. 2000 | 0.659 | 0.667 | -0.012 | 0.665 | 0.595 | 0.108 |
| GMO132[2] | 30 | 100-186 | Brooker et al. 1994 | 0.923 | 0.944 | -0.024 | 0.883 | 0.892 | -0.011 |
| TCH13[2] | 7 | 74-86 | O'Reilly et al. 2000 | 0.919 | 0.889 | 0.033 | 0.924 | 0.919 | 0.006 |

[1] Multiplex 1 (Delghandi et al. 2003)

[2] Multiplex 2 (Dahle et al. 2006; Glover et al. 2010)

Table S3. Tests for deviations from Hardy-Weinberg equilibrium for microsatellite loci used for parentage analysis of Risør cod, calculated separately for each population. No loci deviated significantly from Hardy-Weinberg equilibrium after correction for multiple testing (FDR; α=0.05).

| Locus | P-value $_{(Outer)}$ | P-value $_{(Inner)}$ |
|---|---:|---:|
| GMO35 | 0.124 | 0.331 |
| GMO19 | 0.163 | 0.038 |
| GMO8 | 0.966 | 0.281 |
| TCH11 | 0.196 | 0.444 |
| TCH13 | 0.504 | 0.352 |
| GMO34 | 0.900 | 0.301 |
| GMO2 | 0.509 | 0.074 |
| GMO132 | 0.876 | 0.177 |

Table S4: Identities, collection locations, and ecotype assignments of broodstock and the numbers of offspring included in analysis.

| Parent ID | Location | Ecotype | # of offspring |
|---|---|---|---|
| *Mothers* | | | |
| F03 | Inner | Fjord | 193 |
| F15 | Inner | Fjord | 83 |
| F25 | Inner | Fjord | 168 |
| F31 | Inner | Fjord | 33 |
| F32 | Inner | Fjord | 1 |
| F34 | Inner | Fjord | 15 |
| RIC5060 | Outer | Fjord | 207 |
| RIC5068 | Outer | North Sea | 8 |
| RIC5071 | Outer | Fjord | 115 |
| RIC5078 | Outer | North Sea | 43 |
| *Fathers* | | | |
| F06 | Inner | Fjord | 52 |
| F13 | Inner | Fjord | 7 |
| F14 | Inner | Fjord | 87 |
| F19 | Inner | Fjord | 58 |
| F20 | Inner | Fjord | 117 |
| F21 | Inner | Fjord | 176 |
| F29 | Inner | Fjord | 8 |
| F30 | Inner | Fjord | 32 |
| F33 | Inner | Fjord | 16 |
| F35 | Inner | Fjord | 5 |
| RIC5064 | Outer | Fjord | 60 |
| RIC5076 | Outer | North Sea | 51 |
| RIC5087 | Outer | Fjord | 2 |
| RIC5100 | Outer | North Sea | 195 |
| | | Total | 866 |

Table S5: Genetic cross assignment of larval cod by treatment and replicate.

| Experimental group | | | Genetic cross assignment of larvae | | | | |
|---|---|---|---|---|---|---|---|
| dph | °C | Tank | I×I | I×O | O×I | O×O | Total |
| 0 | 6 | n/a | 7 | 9 | 18 | 6 | 40 |
| 2 | 6 | LT1 | 6 | 10 | 15 | 3 | 34 |
| 2 | 6 | LT2 | 10 | 17 | 12 | 1 | 40 |
| 2 | 6 | LT3 | 16 | 12 | 11 | 0 | 39 |
| 2 | 6 | LT4 | 10 | 16 | 14 | 0 | 40 |
| 2 | 9.5 | IT1 | 10 | 16 | 12 | 2 | 40 |
| 2 | 9.5 | IT2 | 9 | 17 | 12 | 1 | 39 |
| 2 | 9.5 | IT3 | 14 | 17 | 7 | 1 | 39 |
| 2 | 9.5 | IT4 | 12 | 18 | 9 | 1 | 40 |
| 2 | 13 | HT1 | 8 | 20 | 10 | 2 | 40 |
| 2 | 13 | HT2 | 7 | 19 | 10 | 2 | 38 |
| 2 | 13 | HT3 | 12 | 10 | 16 | 1 | 39 |
| 2 | 13 | HT4 | 16 | 10 | 14 | 0 | 40 |
| 28 | 6 | LT2 | 11 | 10 | 14 | 5 | 40 |
| 28 | 6 | LT3 | 13 | 7 | 13 | 7 | 40 |
| 28 | 6 | LT4 | 17 | 9 | 11 | 3 | 40 |
| 28 | 9.5 | IT2 | 10 | 8 | 19 | 3 | 40 |
| 28 | 9.5 | IT3 | 13 | 11 | 12 | 3 | 39 |
| 28 | 9.5 | IT4 | 12 | 7 | 16 | 5 | 40 |
| 28 | 13 | HT1 | 7 | 2 | 28 | 2 | 39 |
| 28 | 13 | HT3 | 7 | 6 | 25 | 2 | 40 |
| 28 | 13 | HT4 | 9 | 6 | 24 | 1 | 40 |
| | | Total | 236 | 257 | 322 | 51 | 866 |

Table S6: Results of a linear mixed effects model for growth for 4 crosses of Atlantic cod, without ecotype included as a random effect. Asterisks denote significance at $\alpha$=0.05.

| Model term | d.f. | Sum of squares | Mean of squares | *F* | *P*-value | |
|---|---|---|---|---|---|---|
| cross | 3 | 0.39 | 0.13 | 0.37 | 0.775 | |
| temperature | 2 | 1.26 | 0.63 | 1.80 | 0.167 | |
| curvature | 1 | 6.36 | 6.36 | 18.11 | <0.001 | * |
| cross×temperature | 6 | 9.94 | 1.66 | 4.72 | <0.001 | * |

| Model term | Variance | SD |
|---|---|---|
| father | 0.04 | 0.20 |
| mother | 0.01 | 0.08 |
| temperature:tank | 0.02 | 0.14 |
| tank | 0.01 | 0.12 |
| residual | 0.35 | 0.59 |

Supplementary Figures

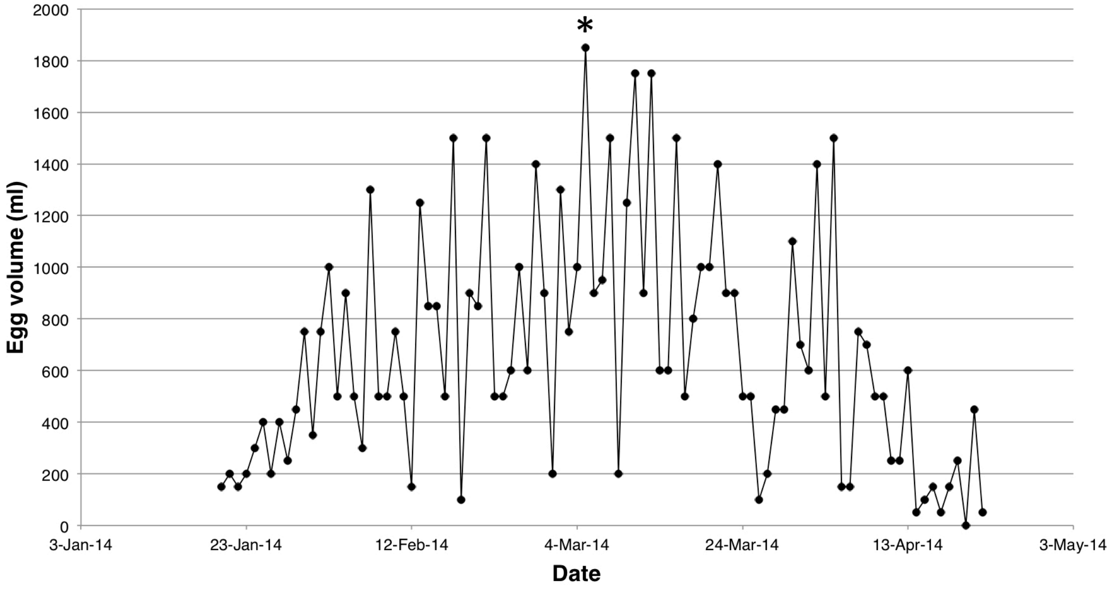


Figure S1: Daily volume of eggs collected throughout the spawning period. The asterisk indicates the eggs that were sampled for the present study.

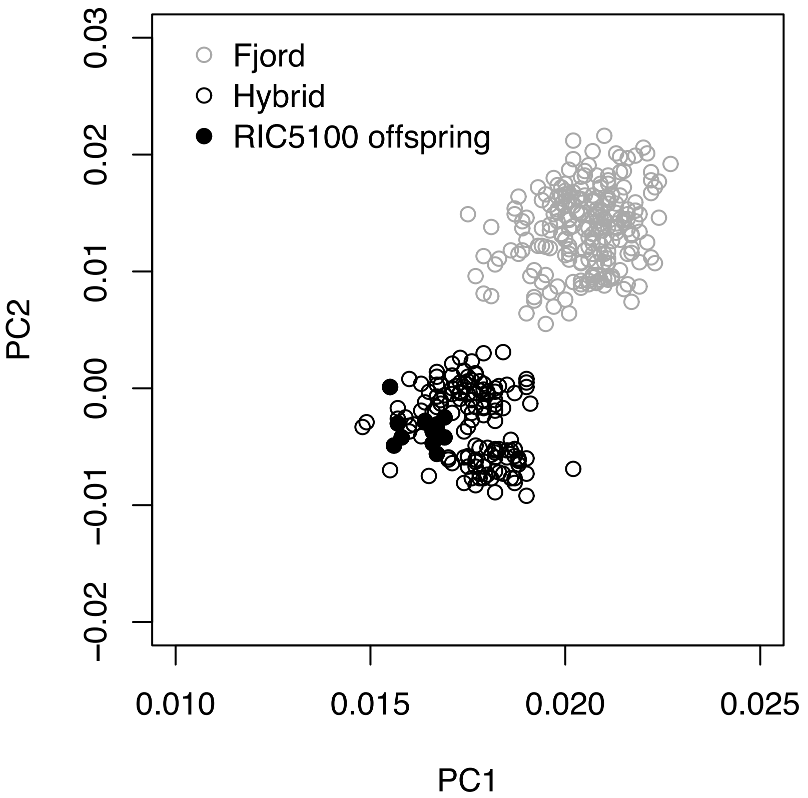


Figure S2: Projected results of a principal component analysis for linkage disequilibrium-purged genome-wide SNPs extracted from reference genome-based transcriptome assemblies of offspring (n=367) from the broodstock in the present study (Oomen, 2019). Principal components were calculated based on a global dataset (n=861; Aqua Genome Project, e.g., see Barth et al., 2019). Fjord and hybrid ecotype classifications are based on parental ecotype assignments from a 26 SNP panel (see Methods). Assignment of RIC5100 was ambiguous based on the 26 SNP panel but classified as North Sea based on the clustering of its offspring produced with fjord mothers with hybrid offspring when using the full set of SNPs.

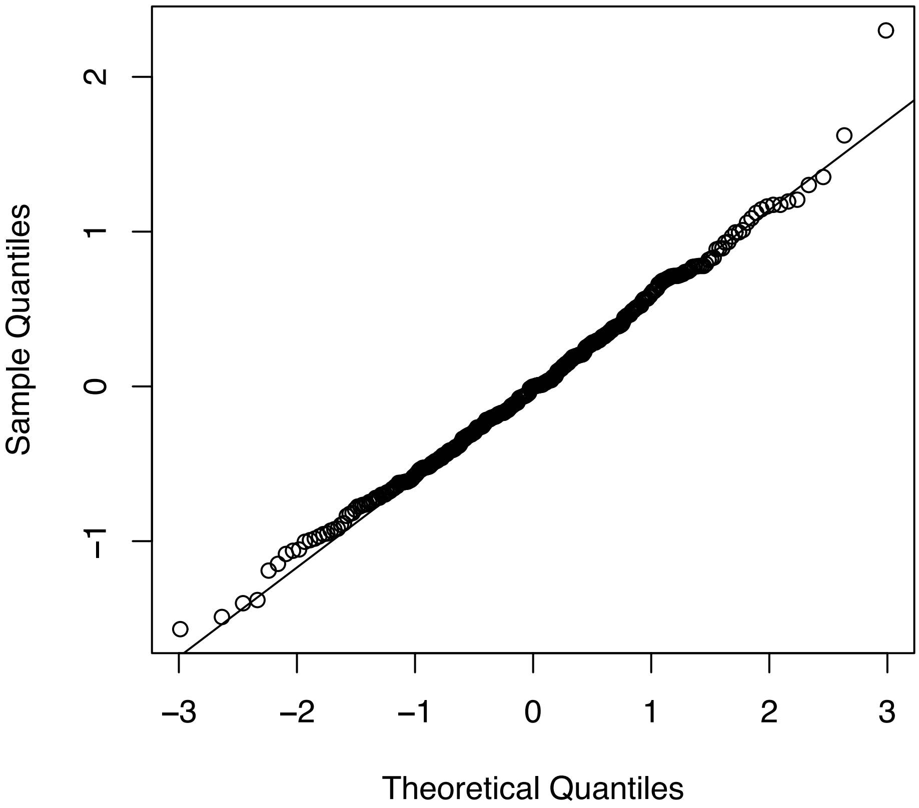


Figure S3: Q-Q norm plot of the residuals from the linear model used in the growth reaction norm analysis.

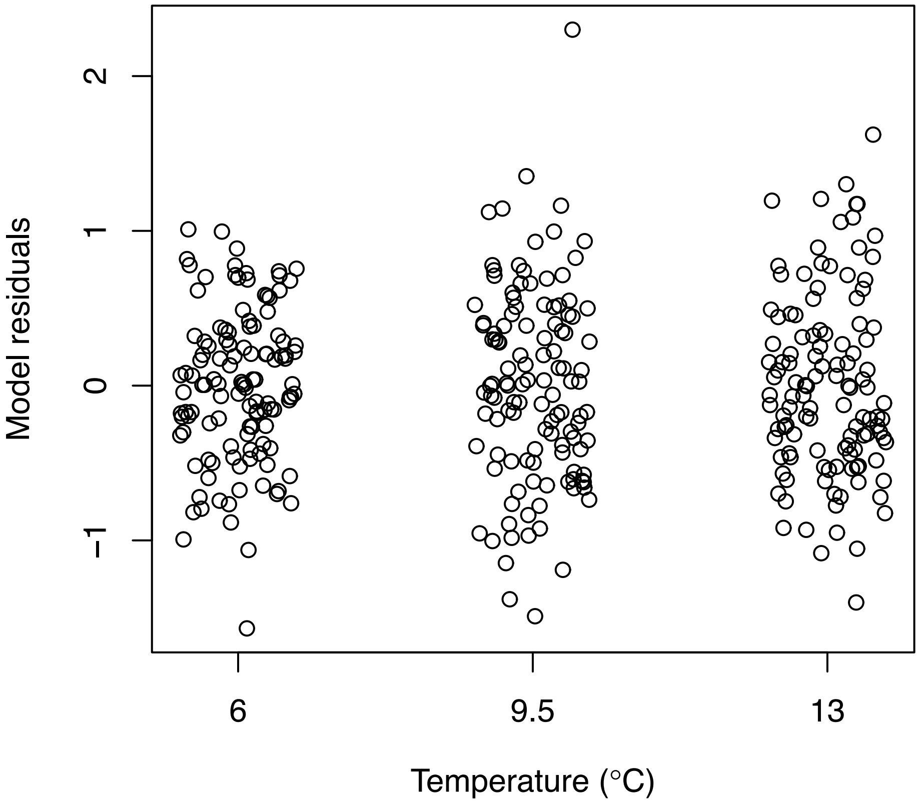


Figure S4: Linear model residuals for each temperature in the growth reaction norm analysis.

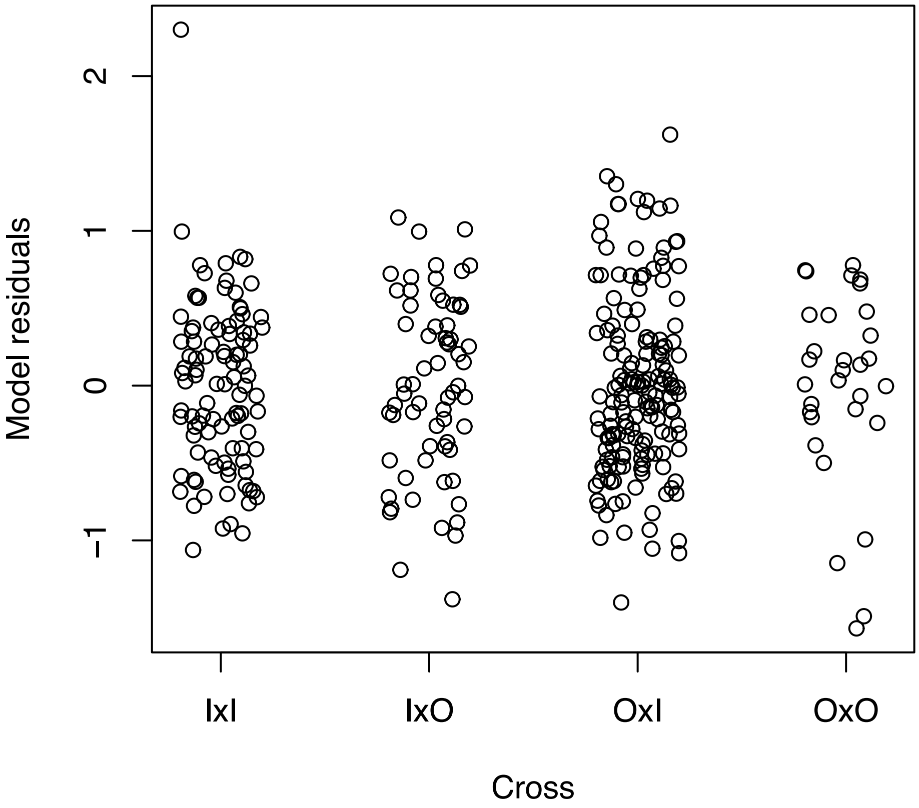


Figure S5: Linear model residuals for each cross in the length-at-day reaction norm analysis.

Figure S6: Thermal reaction norms for larval Atlantic cod growth for four crosses (mother×father: I = inner Risør fjord, O = outer Risør fjord) based on a linear mixed effects model without ecotype included as a random effect. Trendlines represent smoothed conditional means calculated from a loess regression on the model-fitted lengths. Shading represents 95% confidence intervals.

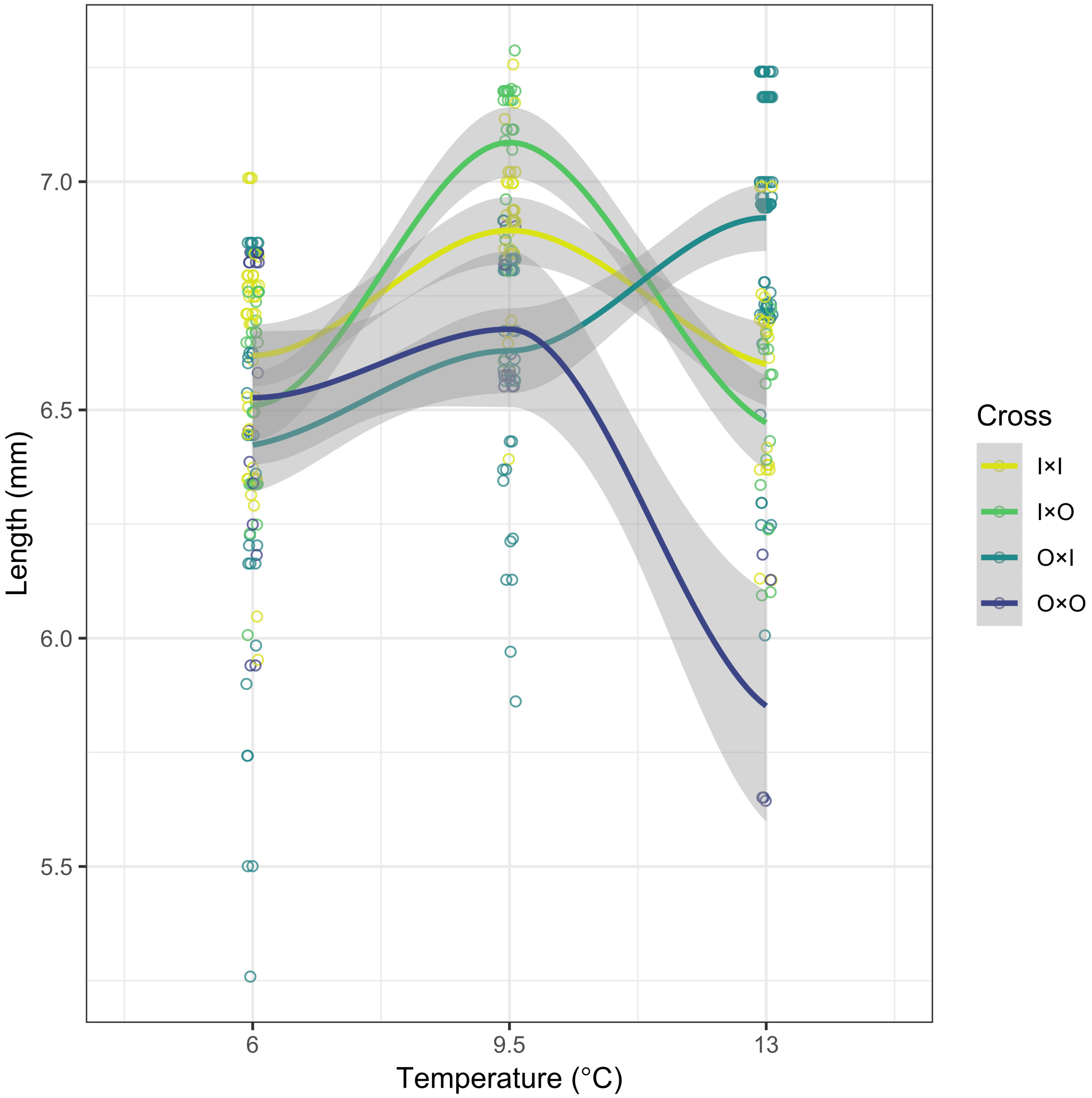

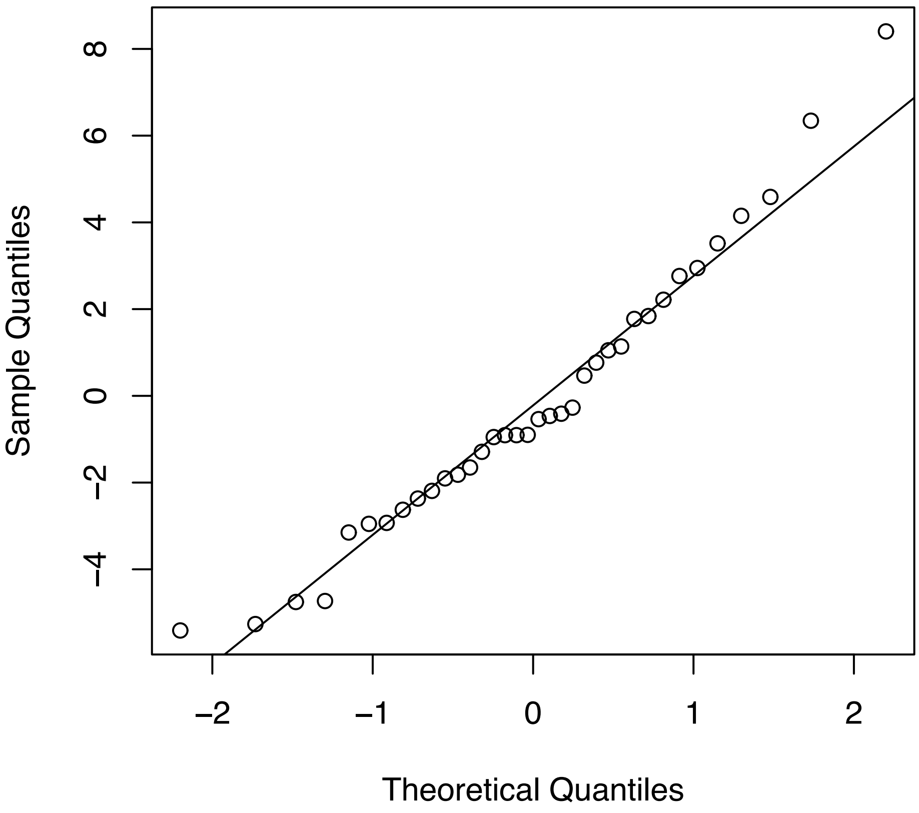


Figure S7: Q-Q norm plot of the residuals from the generalized linear model used in the survival reaction norm analysis.

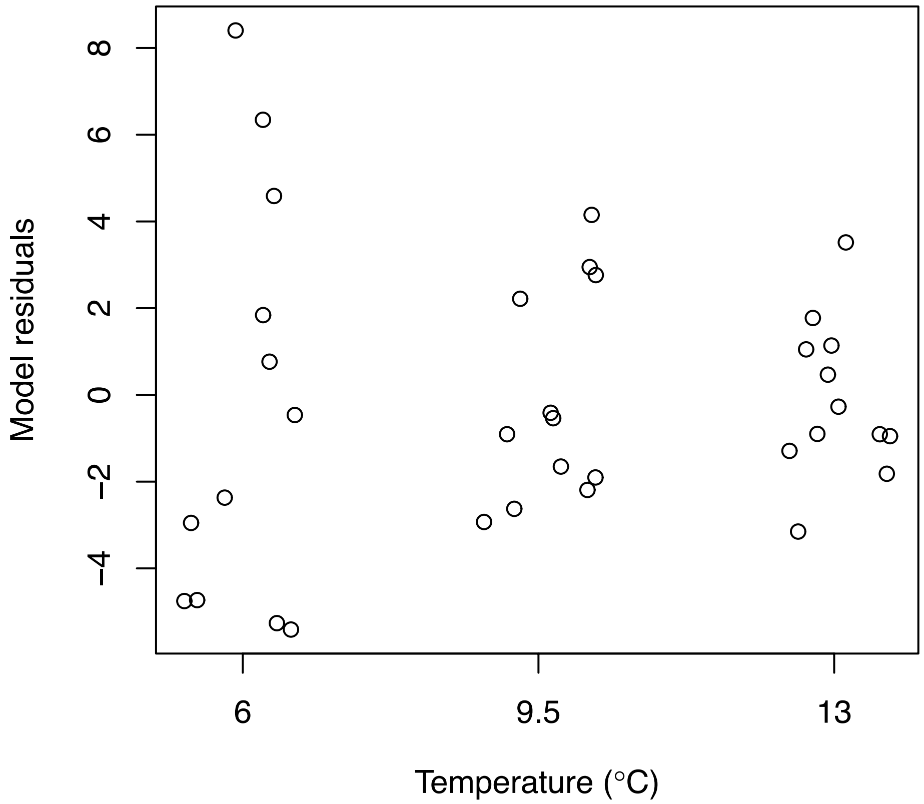


Figure S8: Generalized linear model residuals for each temperature in the survival reaction norm analysis.

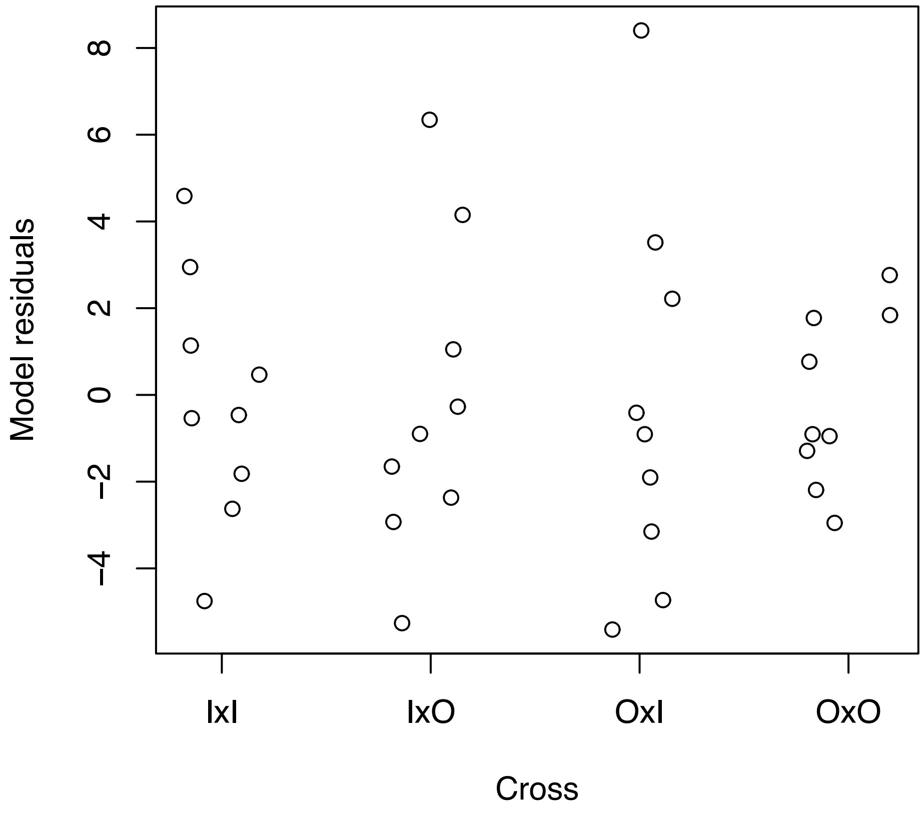


Figure S9: Generalized linear model residuals for each cross in the survival reaction norm analysis.
